# Regulation of Pacemaker Current in the Sinoatrial Node by Zonula Occludens-1

**DOI:** 10.64898/2026.05.19.726291

**Authors:** Rui Zhang, Sarah Teboull, Dexin Chen, Peixin He, Seho Kim, Liang Li, Daniel Adolfo, Terence Gee, Robert S. Ross, Joshua I. Goldhaber

**Affiliations:** Smidt Heart Institute, Cedars-Sinai Health Sciences University, Los Angeles, CA; Department of Medicine, Cardiovascular Medicine Division, University of California, San Diego, La Jolla, CA; Veterans Administration Healthcare, Medicine Section (Cardiology), San Diego, CA

**Keywords:** Zonula Occludens-1, sinoatrial node, arrhythmia, atrioventricular block, bradycardia, arrhythmogenic cardiomyopathy

## Abstract

**BACKGROUND:** In addition to lethal ventricular arrhythmias, arrhythmogenic cardiomyopathy (ACM) is associated with conduction abnormalities, bradycardias, and reduced expression of the scaffolding junctional protein zonula occludens-1 (ZO-1). Reduced ZO-1 expression is also seen in dilated cardiomyopathy, which is far more common than ACM. Conduction abnormalities are likewise a feature of ZO-1 cardiac-specific knockout (ZO-1cKO) mice. However, the role of ZO-1 in sinoatrial node (SAN) automaticity has not been studied.

**OBJECTIVE:** To investigate the role of ZO-1 in SAN automaticity and elucidate the mechanisms by which ZO-1 deficiency leads to SAN dysfunction.

**METHODS:** ZO-1 cardiac-specific knockout (ZO-1cKO) mice were generated by crossing ZO-1 floxed mice with αMHC-nuclear Cre mice. SAN/atrial tissue and isolated SAN cells were examined using optical mapping, single-cell patch clamp, and quantitative PCR techniques to assess functional alterations caused by ZO-1 loss.

**RESULTS:** ZO-1cKO mice exhibited enlarged atria and SAN area compared to control mice, with normal left ventricular function. Electrocardiograms showed sinus bradycardia, sinus pauses and atrioventricular block. Optical mapping revealed a caudal shift in the SAN leading region and reduced intra-atrial conduction velocity in ZO-1cKO mice. Patch-clamp recordings from isolated SAN cells showed reduced spontaneous action potential frequency and diastolic depolarization rate, while voltage-clamp revealed a marked reduction in pacemaker current (I_f_).

**CONCLUSION:** ZO-1 expression is essential for SAN automaticity. Its loss impairs SAN impulse generation by reducing pacemaker current and hampering atrial conduction, leading to bradyarrhythmia, conduction delay and block. These findings help explain impulse generation and conduction abnormalities in ACM and other cardiomyopathies.

## Introduction

The intercalated disk (ID) is a complex cellular structure that connects cardiomyocytes (CMs) both physically and electrically. The canonical description of the ID identifies three types of junctions: adherens junctions (AJ) and desmosomes provide strong attachment between cells while gap junctions (GJ) are critical for synchronous electrical coupling and transmission of small metabolites. Defective function of a range of ID proteins can lead to both myocardial dysfunction and arrhythmias.^1-5^ A key group of proteins located in IDs are those in the zonula occludens (ZO) family.^6, 7^ There are three ZO isoforms: ZO-1, ZO-2 and ZO-3, though only ZO-1 and ZO-2 are expressed in the CMs. Recent studies by our group and others using CM-specific ZO-1 knockout (ZO-1cKO) mouse models have demonstrated that proper expression of ZO-1 in CMs is essential for maintenance of atrioventricular (AV) conduction.^8, 9^ Reduced ZO-1 expression has been identified in human cardiac diseases, including heart failure with reduced ejection fraction (HFrEF), as well as arrhythmogenic cardiomyopathy (ACM).^10-13^ Although ACM is primarily recognized for its strong association with life-threatening ventricular arrhythmias, these patients often also have bradyarrhythmias including sick sinus syndrome. Such arrhythmias are evident in cardiac-specific ZO-1 knockout (ZO-1cKO) mice,^8, 9^ but the mechanism leading to this finding has not been investigated previously. Here we used voltage mapping of an isolated atria/SAN tissue preparation as well as patch-clamping of enzymatically isolated single SAN cells to determine the basis for SAN dysfunction in ZO-1cKO mice. We found a substantial reduction in pacemaker current (I_f_) with associated flattening of diastolic depolarization slope in SAN cell action potentials (APs), along with a clear caudal shift of the leading pacemaker site in the SAN. These results show for the first time that ZO-1 protein plays a critical role in pacemaker activity in the heart and does so by modulating the pacemaker current.

## Methods

The Institutional Animal Care and Use Committee at Cedars-Sinai Medical Center approved all animal protocols, which were performed in accordance with the ethical standards in the Guide for the Care and Use of Laboratory Animals prepared by the Institute of Laboratory Animal Research, National Research Council. Experimental group sizes using both male and female mice were based on previous electrophysiological studies of SAN function in mice.^14, 15^ ZO-1 floxed mice on a C57Bl6 background were crossed with α-Myosin Heavy Chain (α-MHC)-nuclear Cre mice to generate the ZO-1cKO mice.^9, 16^

ZO-1cKO mice were studied using echocardiography and surface electrocardiography (ECG). qPCR was used to assess expression of HCN4 transcripts. To measure membrane voltage and current, we used the patch-clamp technique in enzymatically isolated adult SAN cells. Statistical tests for experiments shown in each figure are reported in the corresponding legend. Data are presented as mean ± standard error of the mean (SEM).

Detailed methods are provided in the online supplement.

## Results

### ZO-1cKO mice have normal contractility but exhibit atrial arrhythmias and conduction block

ZO-1cKO mice are healthy and fertile, without obvious limitations in physical activity. Echocardiography showed normal left ventricular function (Supplemental Figure S1). Surface ECGs on 8- to 12-week-old, lightly anesthetized ZO-1cKO mice revealed variable degrees of AV block, as we described previously,^8^ as well as clear evidence of abnormal atrial rhythm (Figure 1). Overall, loss of ZO-1 was associated with a high degree of variability in atrial rate (PP interval), including frequent pauses (Figure 1A) and prolonged P wave duration compared with control (Figure 1B). We found no change in QRS morphology in the ZO-1cKO vs. paired controls. Notably, AV dissociation and junctional rhythm were prominent in ZO-1cKO mice, along with variable PR and RR intervals (Figure 1C and 1D). Overall, there was a slower ventricular rate in ZO-1cKO mice (438 ± 13 beats per minute [bpm] in ZO-1cKO, N=21, vs. 488 ± 11 bpm in control, N=19, p=0.007, Figure 1E). These findings indicate that ZO-1cKO mice exhibit prominent atrial bradyarrhythmias in addition to our previously identified AV conduction abnormalities.^8^

**Figure 1.**
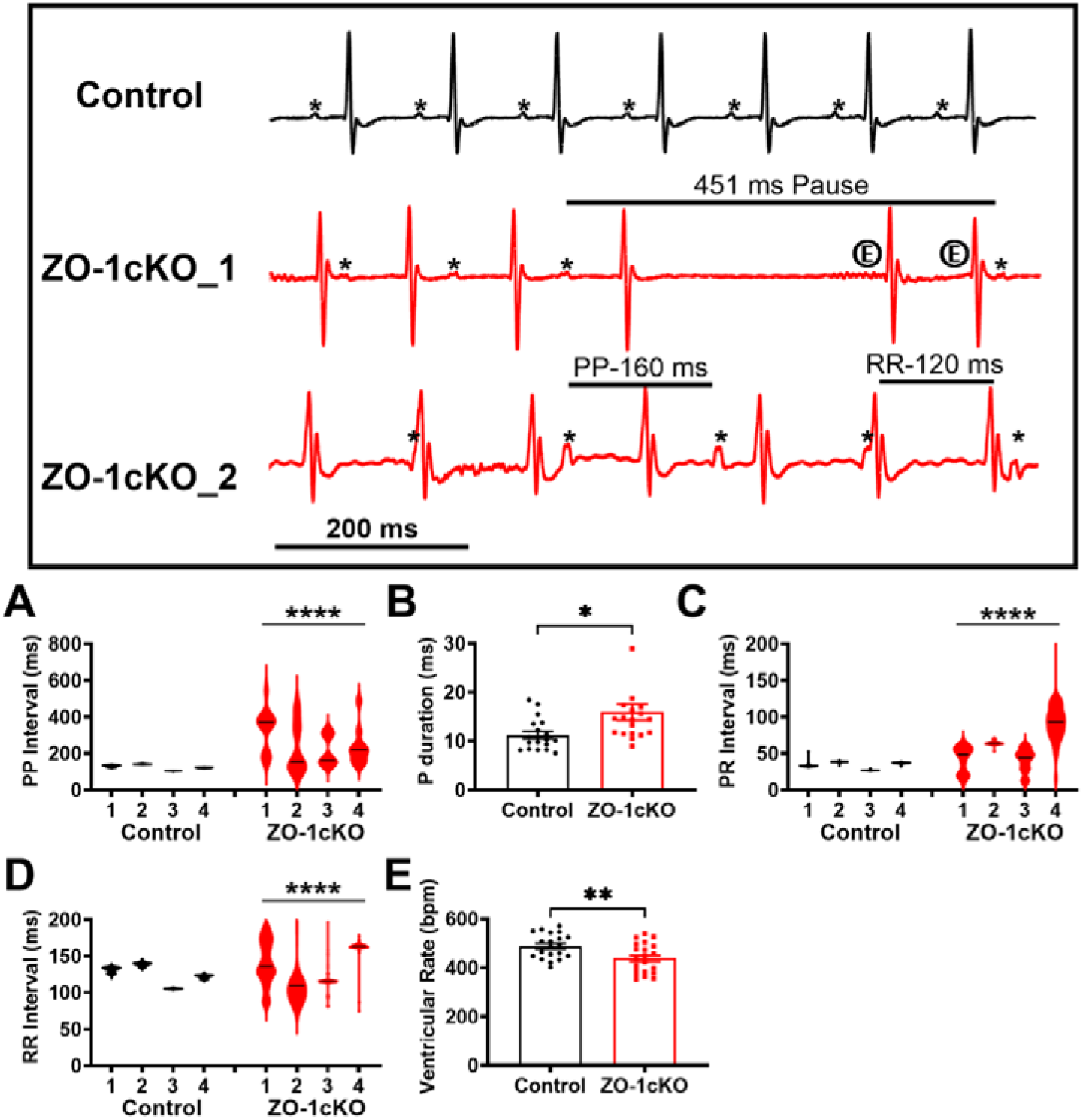
ZO-1cKO mice exhibit prominent atrial bradyarrhythmia as well as AV dissociation and junctional rhythm. **Upper Traces:** Representative surface ECGs recorded in lightly anesthetized 8- to 12-week-old control (*black*) and ZO-1cKO (*red*) mice. Prior to recordings, each mouse was maintained on a heating pad (37° C) and acclimated for 10 min, then data were collected for 1-2 minutes. Control mice showed normal sinus rhythm with a P wave (indicated by *) before each QRS complex, while ZO-1cKO mice exhibited various arrhythmias, most notably frequent sinus pauses (e.g., ZO-1cKO_1 displaying a 451 ms pause and junctional escape beats indicated by circled E) and intermittent AV dissociation indicative of complete heart block and junctional rhythm (ZO-1cKO_2). **A**, Summary data derived from ECG recordings show that PP intervals were highly variable in KO mice compared with control (4 mice surveyed in each group). **B**, P wave duration was greater in KO (19 KO mice vs 18 control). **C**, Increased variability in PR and **D**, RR intervals in KO mice result from intermittent AV block and junctional rhythm with AV dissociation. **E**, Overall, ventricular rate was slower in KO mice (N=21) than control (N= 19). Data normality was evaluated with Shapiro-Wilk test. Data plots are represented as mean ± SEM. Unpaired 2-tailed Student’s test with Welch correction was used to assess the p values by groups.****p<0.0001, **p<0.01, *p<0.05, ns indicates not significant.

### ZO-1cKO mice exhibit SAN/atrial tissue enlargement, abnormal atrial conduction, and a shift in the leading pacemaker site

To investigate the etiology of abnormal atrial rhythm described above, we evaluated atrial and SAN morphology. Microdissection was performed to isolate and separate the right and left atria along with the intact SAN from the ventricles, *en bloc* (Figure 2). The SAN/atrial tissue area was enlarged in KO mice (50.1 ± 1.8 mm^2^ in ZO-1cKO, N=17, vs. 40.3 ± 1.6 mm^2^ in control, N=16, p=0.0003), consistent with atrial size and weights recorded in our previous study.^8^ Next, we used this isolated tissue preparation to perform optical voltage mapping as we have done previously (Figure 3).^14^ In control preparations, APs originated at the classical anatomical location of the SAN, i.e. the junction of the superior vena cava and right atrium. In ZO-1cKO, the leading region of pacemaker activity was shifted caudally and dispersed along the crista terminalis (*dashed line between RA and SAN*, Figure 2). Spontaneous AP rate was reduced and irregular in the ZO-1cKO SAN with intermittent pauses (Figure 3A). In 7 of 10 ZO-1cKO tissues, APs originating in the SAN intermittently failed to excite the left atrium, indicating intra-atrial conduction block (Figure 3, ZO-1cKO_2). Furthermore, conduction velocity from the SAN to the left atrium was significantly reduced in KO compared to control (8.8 ± 3.3 cm/sec in ZO-1cKO, N=11, vs. 26.2 ± 2.3 cm/sec in control, N=11, p=0.0004, Figure 3B). Slowed conduction and chamber enlargement both likely contribute to the prolonged P wave duration observed on surface ECG in KO mice (Figure 1B). Overall, the mapping data indicate that ZO-1cKO mice possess: 1) abnormal SAN pacemaker impulse formation as well as 2) slow and intermittently blocked intra-atrial conduction.

**Figure 2.**
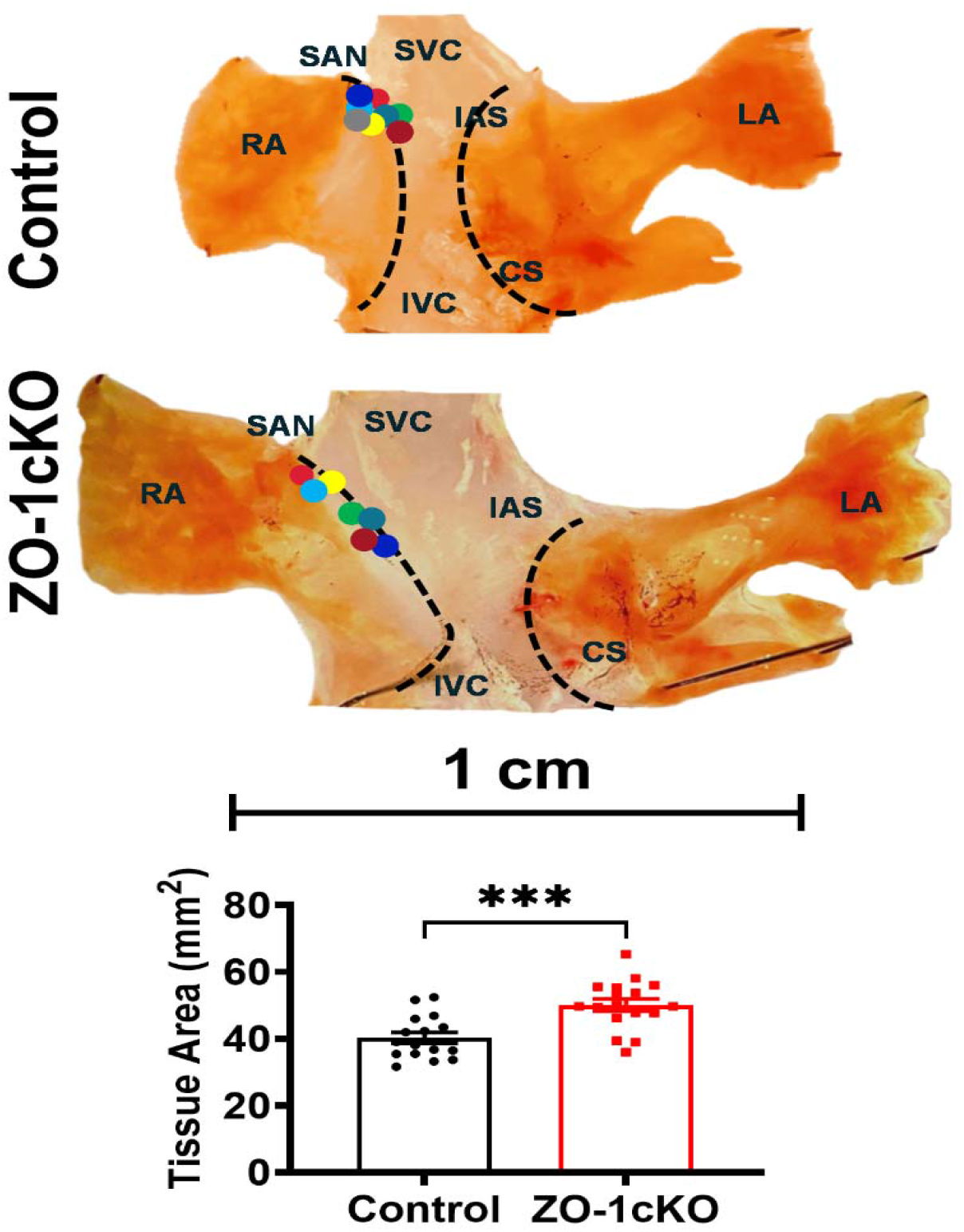
ZO-1cKO mice feature enlarged SAN/atrial tissue area and caudal shift of the leading pacemaker region. Representative images of intact SAN/atrial preparation with demarcations of main features in Control and ZO-1cKO. The sinoatrial node (SAN) is identified between the right atria (RA) and left atria (LA) along the crista terminalis (CT; dashed black line between the RA and SVC/RVC). The SAN/Atria tissue area size was significantly increased in ZO-1cKO vs. control (ImageJ software, NIH, Bethesda, MD). The leading region of pacemaker activity (colored dots) was recorded on SAN tissues loaded with a voltage-sensitive dye (RH237) from 7 KO and 8 control mice and plotted on the representative images above for illustration. In ZO-1cKO, the leading region of pacemaker activity is shifted caudally and dispersed along the CT. Data normality was evaluated with Shapiro-Wilk test. Data plots are represented as mean ± SEM. Unpaired 2-tailed Student’s test with Welch correction was used to assess the p values.****p<0.0001, **p<0.01, *p<0.05, ns indicates not significant.

**Figure 3.**
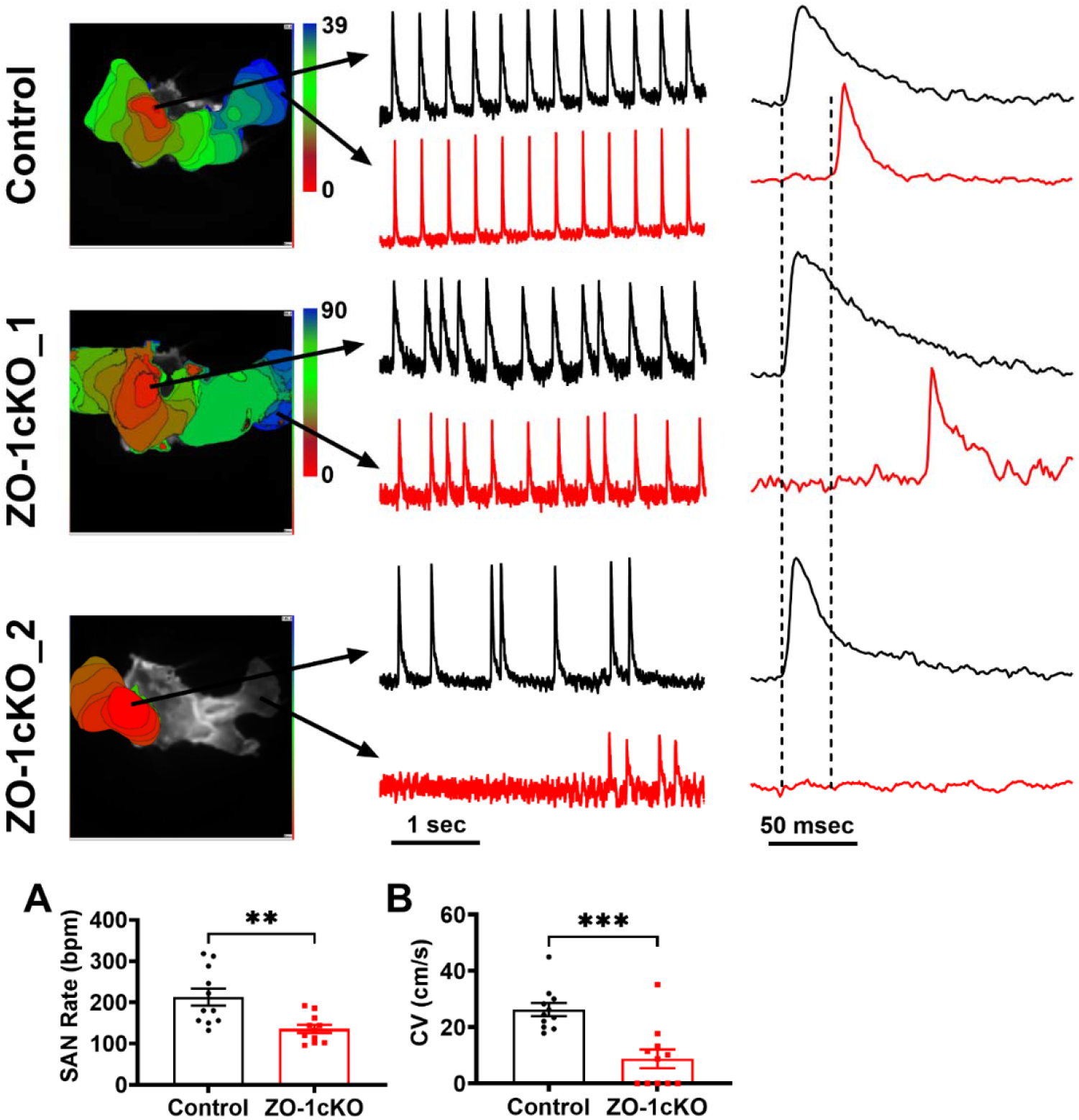
Optical voltage mapping demonstrates impaired conduction velocity and interatrial conduction block in ZO-1cKO mice. **Upper panels:** Representative voltage maps from control and ZO-1cKO SAN/atrial preparations. The earliest activation sites are shown in red and the latest in blue as denoted on the color bar. Sample voltage recordings over time, taken from selected locations in the SAN and LA, are shown to the right of each map, with the SAN traces in *black* and the LA in *red*. Note that in control, there is a regular one-to-one correspondence of the action potentials (APs) in the SAN and LA consistent with normal sinus rhythm. . In the first KO example (ZO-1cKO_1), the APs are irregular. In the second KO example (ZO-1cKO_2), there is no conduction from SAN to LA, and evidence of spontaneous LA activity after a long pause. The voltage traces on the far right are shown on a different time scale to illustrate the delay and block in conduction observed in the KO tissue compared to control. A 50 ms interval is plotted vertically to allow comparison to the typical conduction time for SAN to LA in control mice. **A**, Summary plots show reduced SAN AP rate (bpm) in ZO-1cKO preparations and **B**, slower conduction velocity (CV, cm/s) from the SAN to the LA indicating impaired intra-atrial conduction. N=11 SAN/atrial preparations for each group. Data normality was evaluated with Shapiro-Wilk test. Data plots are represented as mean ± SEM. One-way ANOVA was used to calculate p values. ***p<0.001, **p<0.01.

### Abnormal pacemaker action potentials in isolated ZO-1cKO SAN cells

Next, using the whole cell patch-clamp technique, we recorded paced and spontaneous APs in enzymatically isolated control and KO SAN cells (Figure 4). KO cells exhibited a slower spontaneous AP frequency than control (127 ± 15 AP/min in ZO-1cKO, N=18 cells from 9 mice, vs. 179 ± 13 AP/min in control, N=20 cells from 12 mice, p=0.01, nested t test) without significant differences in maximal diastolic potential (MDP) or peak potential (Figure 4B and 4C). Diastolic depolarization rate (DDR) was slower in KO cells (39.8 ± 5 mV/s in ZO-1cKO vs. 53.5 ± 5 mV/s in control, p=0.04, nested t test, Figure 4E), which is expected in the setting of reduced pacemaker current. AP duration at 50% repolarization (APD50) was reduced in KO compared to control (46.6 ± 6.0 ms in KO vs. 60.1 ± 3.7 ms in control, p<0.05, nested t test) but there was no change in APD90 (Figure 4F). In SAN cells paced in current clamp mode (Figure 5), pacing threshold was increased in KO (0.5 ± 0.03 nA in ZO-1cKO, N=42 cells from 11 mice, vs. 0.4 ± 0.03 nA in control, N=39 cells from 14 mice, p=0.04, Figure 5A). No significant differences were observed in resting membrane potential (RMP), AP amplitude or overshoot (Figure 5B-D). Similar to what we found for spontaneous APs, APD50 was reduced, but APD90 was unchanged (Figure 5E and 5F). Overall, these findings suggest that the prominent atrial bradyarrhythmia observed in ZO-1cKO mice may result, at least in part, from a reduced diastolic depolarization slope and consequent impaired SAN automaticity.

**Figure 4.**
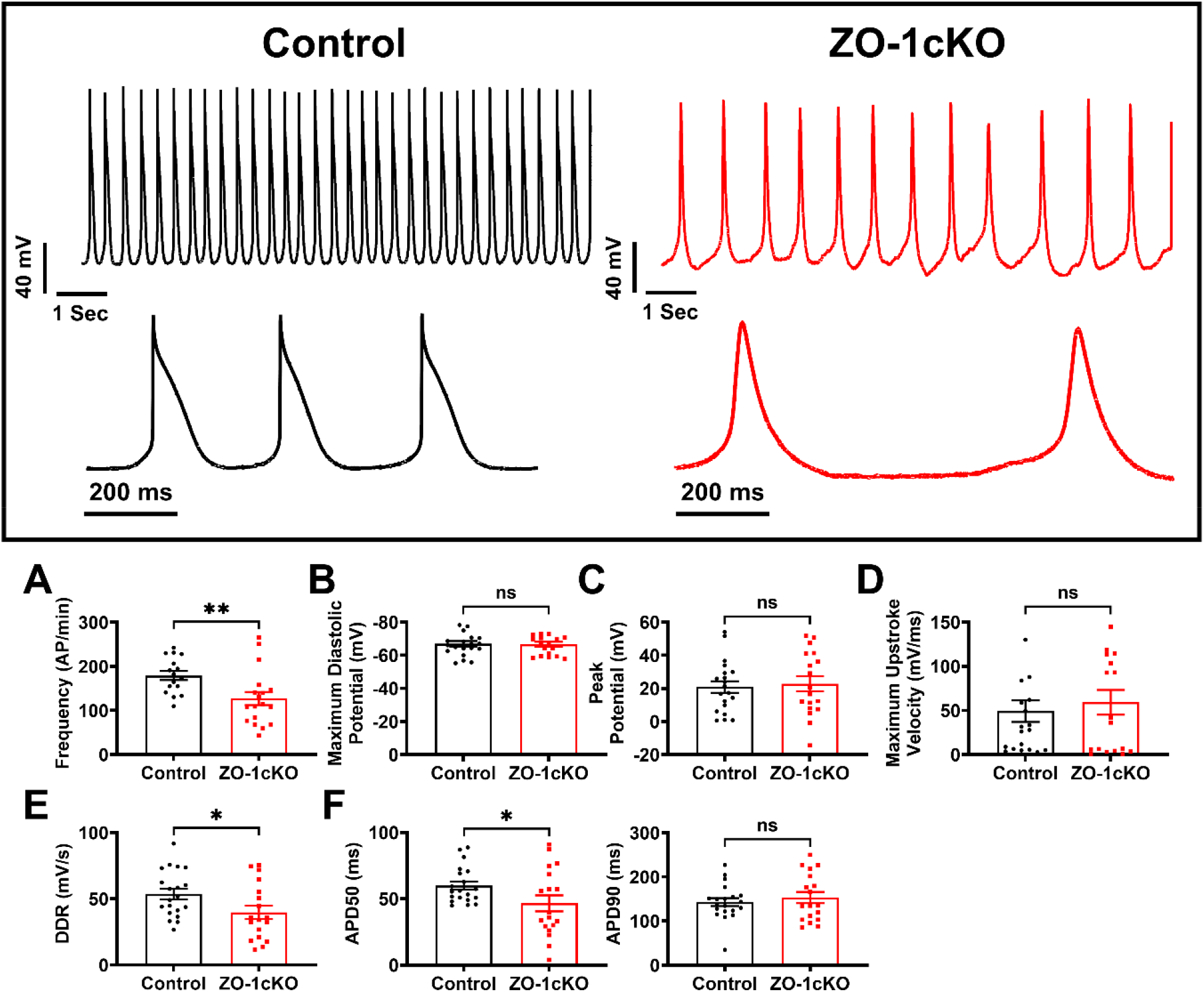
Spontaneous action potentials in patch-clamped ZO-1cKO SAN cells have a reduced diastolic depolarization rate and frequency. **Upper panels:** Representative spontaneous AP traces from control (*black*) and ZO-1cKO (*red*) SAN cells obtained by perforated patch-clamp and displayed on two different time scales. APs occurred at regular intervals in control SAN cells whereas irregular and reduced AP frequency is present in ZO-1cKO. **A**, Summary data show that ZO-1cKO cells exhibited a slower spontaneous firing rate (frequency, AP/min) than control cells, indicating impaired SAN cell automaticity. No significant changes were observed in **B**, maximum diastolic potential, **C**, peak potential or **D**, maximum upstroke velocity. **E**, consistent with our observations, slowed firing rate found in ZO-1cKO SAN cells results from reduced diastolic depolarization rate (DDR) compared to control cells. **F**, AP duration was reduced at 50% repolarization (APD50), but not at 90% (APD90). Data plots were summarized from recordings in 18 cells from 9 ZO-1 cKO mice and 20 cells from 12 control mice. Nested t test was used to calculate p values. **p<0.01, *p<0.05, ns indicates not significant.

**Figure 5.**
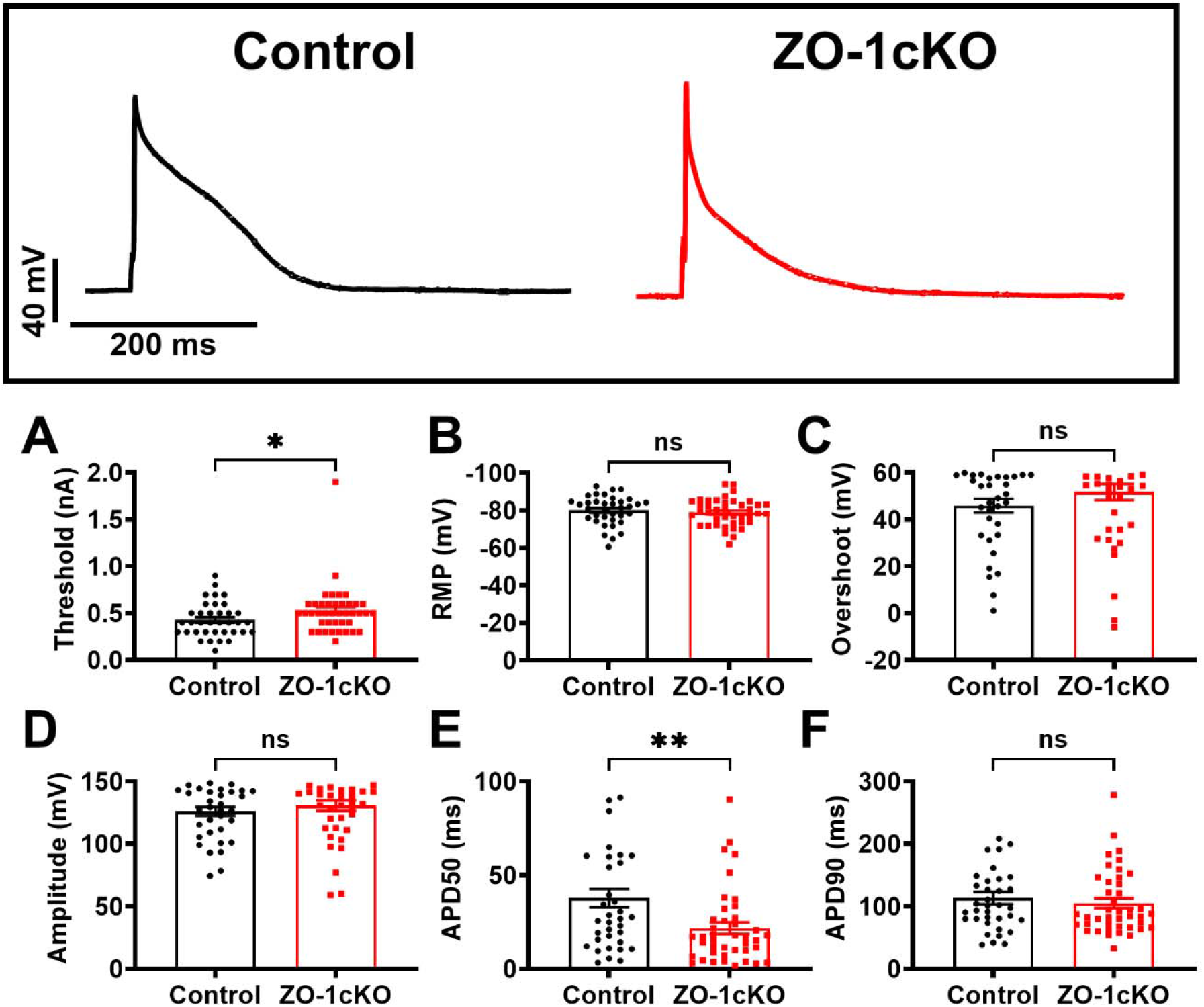
Stimulated AP recordings from patch-clamped isolated SAN cells show increased threshold and reduced APD50 in ZO-1 cKO SAN cells. **Upper panels** show representative traces of stimulated APs recorded in SAN cells isolated from control (*black*) and ZO-1cKO (*red*) mice. **A**, ZO-1cKO paced SAN cells exhibited an increased pacing threshold compared to control cells, but no significant differences were observed in **B**, resting membrane potential (RMP), **C**, overshoot and **D**, AP amplitude. Similar to spontaneous AP recordings, ZO-1cKO SAN cells had reduced APD50 compared to control (**E**) but no changes in APD90 (**F**). Data plots were summarized from recordings in 42 cells from 11 ZO-1cKO mice and 39 cells from 14 control mice. Plots are represented as mean ± SEM. Nested t test was used to calculate p values. **p<0.01, *p<0.05.

### ZO-1cKO SAN cells have reduced pacemaker current (I_f_)

To determine the mechanism underlying the reduced diastolic depolarization slope and spontaneous AP rate of patch-clamped SAN cells in ZO-1cKO mice, we recorded pacemaker current (funny current, I_f_) by applying 3s hyperpolarizing voltage steps from –50 to –160 mV in 10 mV increments from a holding of potential of –50 mV (Figure 6A).^17^ Current-voltage plots of I_f_ in KO were significantly decreased (Figure 6B) without change in conductance (Figure 6C). At -140 mV I_f_ was decreased by 63.8% (N=38 cells from 10 ZO-1cKO mice compared to N=41 cells from 10 control mice, Figure 6D). Double exponential fits of full-activated currents (at –140 mV) revealed similar activation kinetics in KO cells compared to control (Figure 6E). Given that whole cell current, *I*=*iNP*_o_, our findings suggest that I_f_ is reduced primarily because of a lower number (*N*) of HCN4 channels at the membrane, rather than reduction in single-channel current (*i*) or channel open probability (*P*_o_). Consistent with this interpretation, HCN4 gene expression was also reduced 53.1 ± 6.2% in ZO-1cKO SAN/atrial tissue (Figure 6G).

**Figure 6.**
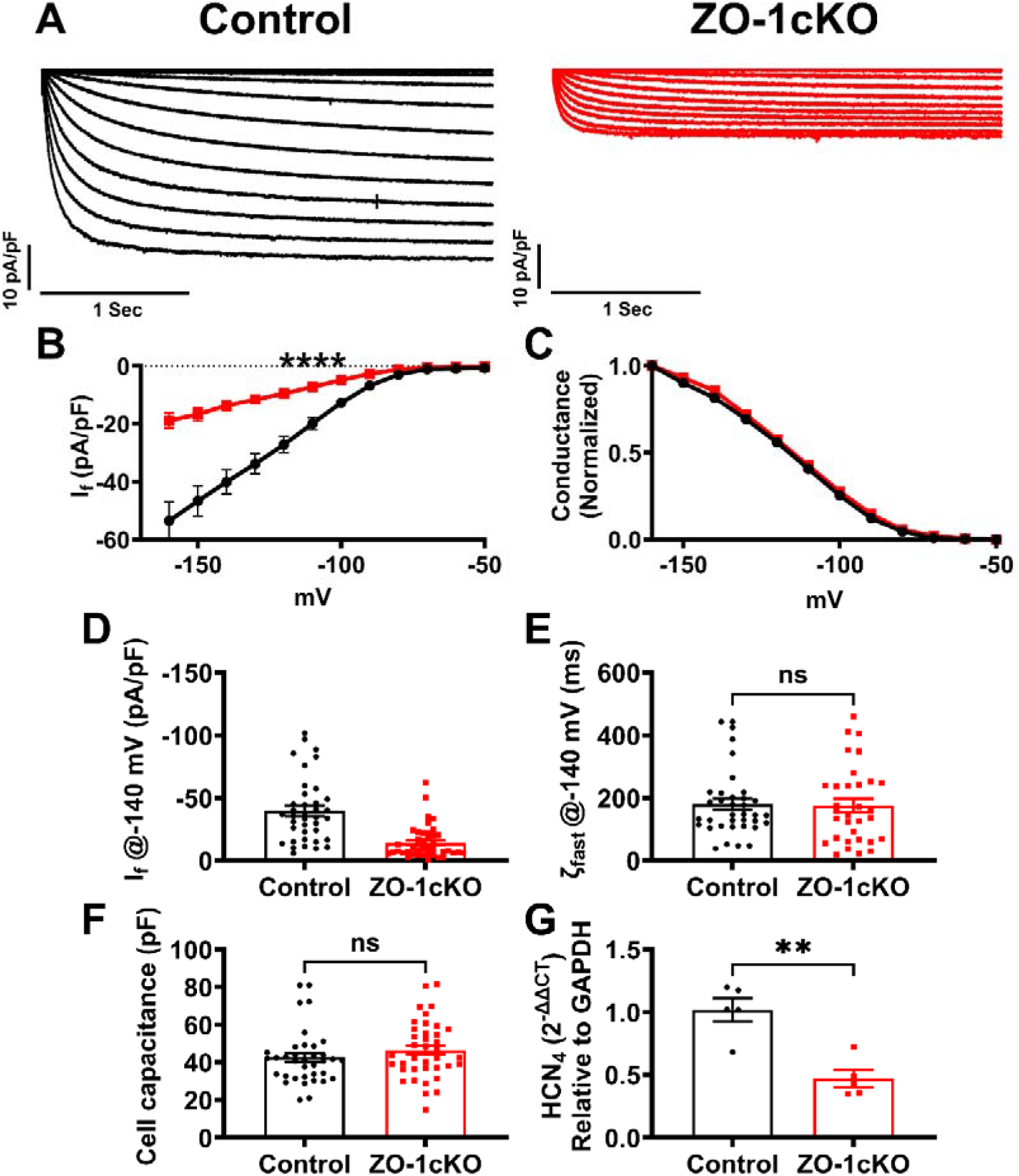
Pacemaker current (I_f_) and HCN4 expression are reduced in ZO-1cKO SAN. **A**, Representative I_f_ current families (normalized to cell capacitance) from control (*black*) and ZO-1cKO (*red*) SAN cells. I_f_ was recorded by applying 3 s hyperpolarizing voltage steps from –50 to –160 mV in 10 mV increments from a holding of potential of –50 mV. **B**, summary plot showing reduction in mean I_f_ current-voltage relationship in ZO-1cKO SAN cells. **C**, there was no change in normalized conductance. **D**, I_f_ density at -140 mV was found significantly reduced in ZO-1 cKO SAN cells compared to control. No significant differences were observed in **E**, the rapid phase of current decay (ζ_fast_) and **F**, cell capacitance. Data plots were summarized from recordings in 38 cells from 10 ZO-1 cKO mice and 41 cells from 10 control mice. **G**, HCN4 gene expression normalized to GAPDH was found significantly reduced in ZO-1cKO SAN/atrial tissue (N=5 for each group). Data plots were analyzed by two-way ANOVA or two-tailed Mann-Whitney test and presented as mean ± SEM. ****p<0.0001, **p<0.01, *p<0.05, ns indicates not significant.

## Discussion

In this study, we used a cardiac myocyte-specific KO mouse model of the zonula occludens protein ZO-1 to specifically investigate the mechanistic role of ZO-1 in SAN function. ZO-1cKO mice not only exhibit the AV conduction abnormalities described previously, but also feature sinus bradycardia and pauses, accelerated junctional rhythm and AV dissociation (Figure 1). Optical voltage mapping revealed that ZO-1 deficiency shifts the leading pacemaker region caudally to less active subsidiary pacemakers and slows intra-atrial conduction (Figure 2). Critically, single SAN cell patch-clamp recordings reveal markedly reduced pacemaker current (I_f_), resulting in defective impulse generation (Figure 6). These results demonstrate that ZO-1 plays a critical role in SAN pacemaker function and atrial conduction. This could have important implications for human diseases, particularly dilated and arrhythmogenic cardiomyopathies where ZO-1 deficiency has been described.

### Reduction in HCN4 channel current and expression

Pacemaker automaticity in the SAN depends upon the coupled activity of the membrane clock, dominated by depolarizing currents generated by HCN4, Ca_v_1.3 and Ca_v_3.1 channels,^18^ and the calcium (Ca) clock, where spontaneous release of sarcoplasmic reticulum (SR) Ca depolarizes the cell upon exit through the electrogenic Na/Ca exchanger.^8, 17, 19 20^ A third coupled oscillator, also known as the mechanics clock,^21^ has been proposed as a mechanism underlying the observation that stretching pacemaker tissue can generate spontaneous beating, and that right atrial stretch in response to a fluid bolus can increase heart rate in anesthetized animals.^22^ Many mechanosensitive ion channels have been identified in the SAN, including Piezo1 and 2, and it is well known that stretch-induced activation of reactive oxygen species (ROS) signaling, also known as X-ROS, in cardiac myocytes can influence intracellular Ca release.^23^ As a scaffolding protein located in the ID, it is reasonable to consider ZO-1 as a potential contributor to the regulation of the mechanics clock, but we did not investigate that mechanism here.

We found that SAN myocytes from ZO-1cKO mice show a profound reduction in HCN4-mediated I_f_, the major component of the membrane clock, and a corresponding reduction in HCN4 gene expression (Figure 6). In the absence of a shift in current-voltage dependence, there is no evidence of altered regulation of expressed HCN4 channels. These findings suggest a fundamental relationship between ZO-1 expression and HCN4 transcription. To the best of our knowledge, there is no known association of ZO-1 and HCN4 gene expression. Transcription factors active during development (e.g., TBX18, SHOX2) are critical for promoting HCN4 gene expression and supporting the pacemaker cell phenotype.^24^ Secondary reduction in HCN4 in transgenic models has been reported previously, most notably in the setting of mitochondrial Trx2/Txn2 conditional deletion. This reduction is hypothesized to be caused by increased expression of HDAC4, mediated by reactive oxygen species.^25^ Unlike reductions in proteins such as Cx45 in KO models of ZO-1 or coxsackievirus and adenovirus receptor (CAR) described previously,^8, 26^ there is no known direct physical linkage of ZO-1 and HCN4. Further study is needed to determine the mechanism underlying reduced I_f_ in this KO model.

### Sinus bradycardia and arrhythmia

Reduced HCN4 expression and I_f_ explain the reduction in AP diastolic depolarization rate (Figure 4), which accounts for the slowing of SAN pacemaker activity measured as PP interval on ECG (Figure 1). We also observed a caudal shift in the leading region of pacemaker activity in the SAN (Figure 2). It is well established in multiple mammalian species, that the dominant pacemaker regions and earliest activation sites within the SAN can shift to different locations depending on disease states and autonomic activity that can be mimicked through the use of muscarinic and β-adrenergic agonists.^27-30^ While the caudal shift in ZO-1cKO cells could indicate that I_f_ reduction is more pronounced in the superior region of the SAN, thereby enabling subsidiary pacemaker sites along the crista terminalis, it is also possible that tissue remodeling in the KO distorts the anatomy and shifts the most active SAN pacemaker cells to a more inferior location. One limitation of voltage mapping is that we cannot necessarily perceive depolarization of one or a few SAN cells. To be detected, a large enough area would have to be depolarized, which not only depends upon I_f_ function, but also intercellular communication. We found previously that Cx45, the dominant connexin in the SAN, is reduced in this KO model.^8^ Not only could this alter the location of the leading region as suggested above but could readily account for the reduced intra-atrial conduction velocity we observed in these mice. (Figure 3B). Notably, increased fibrosis has not been observed in ZO-1cKO mice.^8^

Our prior study using this mouse model, as well as a second one that used a varied model to reduce ZO-1 expression in cardiomyocytes,^8, 9^ found that ZO-1 is essential for proper AV node conduction. Our prior work^8^ also noted a reduction in atrial rate but did not investigate the underlying mechanism. The study by Dai et al^9^ used a tamoxifen-induced ZO-1 cardiac myocyte KO model, which excised the TJP1 gene only in the adult mouse heart. Notably, that study found there was no change in atrial rate. We speculate that the fundamental differences in the two models, likely the timing of gene excision, accounts for this important difference. The implication is that embryonic onset of ZO-1 deletion, as occurs in the ZO-1cKO model we use here, may be necessary for disruption of SAN activity by reduced expression of I_f_. However, both models feature AV block, showing the clear importance of proper ZO-1 expression to preserve normal conduction, regardless of the timing of ZO-1 loss.

### Action Potential Changes

Aside from reduction in diastolic depolarization rate, we found that APs in isolated SAN cells from our ZO-1cKO mice feature more rapid repolarization, with reductions in APD50 (Figure 4F,5E). AP duration is determined by the balance of ion influx and efflux, with reductions typically caused by either a reduction in depolarizing inward Ca current (e.g., I_Ca,L_), or an increase in repolarizing outward K current (e.g. I_TO_). Notably, we did not detect a change in maximum upstroke velocity (Figure 4D), which in SAN cells is dominated by I_Ca_. This finding argues against reduced I_Ca_ in this model, and points towards increased repolarizing currents as the most likely cause of shortened APD50. In paced, patch-clamped KO cells, we observed an increase in the pacing threshold, i.e. the amount of current injection needed to stimulate an AP (Figure 5A). Threshold is determined by many different factors but decreased I_f_, decreased I_Ca_ and increased I_K_ are likely to increase threshold. In future studies it will be important to selectively assess these currents using voltage clamp.

### Relevance to human disease

Ventricular and atrial tachyarrhythmias accompany virtually all forms of cardiomyopathy. While lethal ventricular arrhythmias have always been a major concern for cardiomyopathy patients, up to half of these patients actually die from sinus bradycardia, asystole or pulseless electrical activity.^31, 32^ Rhythm disturbances specific to the SAN, including sinus bradycardia, are common in heart failure.^33^ Atrial fibrillation and sinus node disease frequently coexist, suggesting causation or at least common pathophysiology.^34^ Consistent with that concept, it is well established that the prevalence of atrial fibrillation is considerably higher in patients with heart failure compared to the general population, particularly in HFpEF.^34^ ACM is notable for risk of sudden death due to ventricular tachycardia, but these patients also have a higher incidence of sinus node disease than the normal population.^35^ The etiology of bradyarrhythmias is multifactorial but can be generally categorized as defects in impulse formation and/or conduction, often involving abnormal behavior of ion channels or disrupted cell-to-cell connections, and frequently associated with atrial remodeling. Disorders of cell-to-cell communication/connections are well-described causes of conduction abnormalities.^36^ In this regard, abnormalities of desmoplakin and connexins have been investigated.^37, 38^ Pathogenic variants and altered expression of ZO-1 have been observed in ACM and heart failure.^12^ The SAN pacing and atrial conduction abnormalities we describe here are consistent with abnormalities observed in the various cardiomyopathies described above. This suggests that further study is warranted to understand the detailed mechanism in search of a therapeutic intervention.

### Limitations

Several limitations should be acknowledged when interpreting our findings. First, although we observed reduced I_f_ in ZO-1cKO SAN cells, we did not assess other currents that contribute to SAN automaticity. In particular, L-type and T-type calcium currents (I_Ca,L_ and I_Ca,T_) through Ca_v_1.3 and Ca_v_3.1 channels, which have been implicated in diastolic depolarization,^39, 40^ were neither functionally assessed nor quantified at the expression level. As such, it remains unclear whether ZO-1 deletion also affects the broader membrane clock machinery or only specific components. We also did not investigate I_Na_ in SAN cells from ZO-1cKO mice. Although not usually considered a key component of the pacemaker machinery, because it is absent in central SAN cells, several studies have identified I_Na_ and Na_V_1.5 in more peripheral SAN cells, where it is thought to help propagate the pacemaker impulse from the central SAN to the surrounding atrial tissue. Prior studies of ZO-1 KO have found a reduction in Na_V_1.5 in ventricle, so it is reasonable to suspect that this channel is also reduced in SAN cells, which could lead to sinus node exit block.

SAN automaticity is governed not only by the membrane clock but also by a Ca clock with multiple regulatory components, including SERCA, RYR2 and NCX,^17, 41, 42^ We have not yet examined the performance of the Ca clock in this model, which could reveal additional defects beyond those we found in the membrane clock. Despite these limitations, the very large reduction in I_f_, the dominant pacemaker current in SAN,^43, 44^ is sufficient to account for the abnormal automaticity we observe in this mouse model. As noted above, the mechanics clock also adds an additional level of complexity to SAN function. Tissue stretch can lead to alterations in automaticity, and while we sought to avoid overstretching of the SAN tissue preparation during experiments, it is possible that we did so inadvertently. However, this is unlikely given precautions we take against this, namely applying blebbistatin to reduce contraction, and use of consistent technique with each tissue preparation. Related to this concern is the possibility that the mechanics clock is impaired by ZO-1 ablation, which could contribute to the SAN dysfunction observed in this study. Future studies will be required to address this issue.

Finally, we know from our prior studies that ZO-2 is increased in ventricular myocytes from ZO-1cKO mice. We did not measure ZO-2 SAN expression in the ZO-1cKO SAN cells, but based on the ventricular cell data, it is reasonable to hypothesize that ZO-2 is increased in the SAN. It is unknown if ZO-2 overexpression aggravates or mitigates SAN dysfunction in the absence of ZO-1. Further studies will be necessary to assess the effect of ZO-2 loss on SAN function.

## Conclusion

Our findings establish ZO-1 as a critical determinant of SAN automaticity. Loss of ZO-1 disrupts expression of the SAN primary pacemaker channel (HCN_4_) resulting in reduced pacemaker current (I_f_) dysfunctional pacing activity, and impaired atrial conduction. While not studied here, the intracellular “Ca clock” driven by rhythmic spontaneous Ca release from the sarcoplasmic reticulum (SR)^21, 45, 46^ may also contribute to SAN automaticity. It will be important to examine this mechanism, and the mechanics clock, in future studies.

## Abbreviations

AP: Action Potential
AV: atrioventricular
DDR: Diastolic Depolarization Rate
GAPDH: Glyceraldehyde 3-phosphate dehydrogenase
HCN4: Hyperpolarization-Activated Cyclic Nucleotide-Gated Potassium Channel
I_f_: Funny current (pacemaker current)
KO: knockout
SAN: Sinoatrial node
ZO-1: Zonula Occludens-1
ZO-1cKO: Cardiac-specific ZO-1 knockout

## Declaration of generative AI and AI-assisted technologies in manuscript preparation

Generative AI was not used in the preparation of this manuscript.

