## Supplementary material for "Regulation of Pacemaker Current in the Sinoatrial Node by Zonula Occludens-1": Online Supplement

### Detailed Methods

#### Generation of ZO-1cKO mice

The Institutional Animal Care and Use Committee at Cedars-Sinai Medical Center approved all animal experiments, which conformed to the National Institutes of Health Guide for the Care and Use of Laboratory Animals. ZO-1 (exon 3) floxed mice (ZO-1$^{fx/fx}$) on a C57Bl6 background were crossed with ZO-1$^{fx/fx}$ α-Myosin Heavy Chain (α-MHC)-nuclear Cre (nCre) mice to produce cardiomyocyte-specific ZO-1 knockout mice (ZO-1cKO)[1]. All animals were housed in the Cedars-Sinai Department of Comparative Medicine animal facility, which is fully accredited by the Association for the Assessment and Accreditation of Laboratory Animal Care (AAALAC). Both male and female mice were used in all experiments. Consistent with prior studies, ZO-1 gene expression was reduced by 59.0 ± 7.2 percent in ZO-1 cKO mice compared with control (N=5 for each group).

#### Echocardiography

We anesthetized eight- to twelve-week-old mice using 3% isoflurane for induction and 1.5% isoflurane for maintenance. The animals were placed in a supine position on a heated platform (37$^{o}$C) for vital sign monitoring, and then transthoracic echocardiography was performed with continuous electrocardiographic (ECG) monitoring using a Vevo 3100 machine (VisualSonics, Toronto, Canada). We measured ejection fraction (EF) and wall thickness, calculated from M-mode images obtained in the parasternal short-axis view at the level of the papillary muscles. Three separate measurements were averaged for each animal.

#### Electrocardiography

We performed electrocardiograms (ECGs) during echocardiography by placing three 12 mm needle electrodes (29 gauge, MLA1213, ADInstruments, Dunedin, New Zealand) subcutaneously in the standard lead II configuration. Rhythm strips were recorded using amplifying and recording equipment (PowerLab 8/30, equipped with an Animal Bio Amp and LabChart7 Pro ECG analysis software - ADInstruments,

Dunedin, New Zealand). Each channel was amplified and sampled at a rate of 120 Hz with a 2 mV range, and a high-pass filter setting of 0.03 seconds.

**Intact SAN/Atrial Preparation.**

To isolate intact SAN/atrial tissue for experimentation, mice were heparinized (300 U i.p.), anesthetized deeply with 33% isoflurane, then thoracotomy was performed and hearts excised. The atria and SAN were separated *en bloc* from the ventricles and pinned to the bottom of an optical chamber (Fluorodish, FD35PDL-100; WPI, Sarasota, FL) coated with approximately 5 mm of clear silicone elastomer (Sylgard 184, Dow Silicones Corporation, Midland, MI) to maintain the SAN in a flat plane. The chamber contained heparinized (10 U/mL) modified Tyrodes solution heated to 36 °C and containing (in mM): 136 NaCl, 5.4 KCl, 10 HEPES, 0.33 $NaH_2PO_4$, 10 Dextrose, 1 $MgCl_2$, and 1.8 $CaCl_2$, pH adjusted to 7.4. This solution also served as the control solution for all experiments. Using a stereomicroscope (SZX16; Olympus, Tokyo, Japan) with 7x magnification, the tissue was transilluminated and directly visualized. The SAN region was identified using the borders of the superior and inferior vena cavae, the crista terminalis, and the interatrial septum as landmarks, as we described previously [2, 3]. Images photos were obtained on the stereomicroscope described above at 10x magnification using a standard iPhone 15 Pro Max (Apple Inc, Cupertino, CA) attached directly to the right eyepiece (optical zoom set to 1). With the aid of a physical scale bar in the photo, the area of the SAN/atria in $mm^2$ was calculated using ImageJ (NIH, Bethesda, MD) software[4].

**Optical Voltage Mapping**

Voltage changes in the SAN/atrial tissue were assessed by optical mapping. The tissue preparation described above was immersed in Tyrodes solution containing the voltage-sensitive dye RH237 ([10 μM]; Biotium, Fremont, CA) for 30 min at 34–36 °C on a stirring hotplate. Continuous agitation and oxygenation were maintained during dye loading. The tissue was subsequently washed in dye-free Tyrodes solution for 15 min at 34–36 °C and then perfused with 1.8 mM Ca Tyrodes containing 5 μM blebistatin (Tocris

Bioscience, Ellisville, MO) to eliminate motion during imaging. Optical mapping was performed at 1000 frames/s (1 ms/frame) using a MiCAM05 Ultima-L CMOS camera (100 × 100 pixels; SciMedia, Costa Mesa, CA) mounted on a twin high-magnification tandem (THT) microscope with a 0.63× objective (16 × 16 mm field of view). Excitation was provided by a 150-W halogen light source with a built-in shutter (SciMedia, Costa Mesa, CA) and a filter set which included a 520/35-nm excitation filter, 560-nm dichroic mirror, and 580 ± 40-nm long-pass emission filter. Recordings were limited to 10 s to minimize phototoxicity, and data were analyzed using BrainVision Analyzer software (v16.04.20 (BrainVision Inc. Tokyo Japan). Recordings were spatially filtered using 3x3 pixel matrix, and activation maps were generated by calculating 50% of the maximum amplitude. The earliest activation site was identified as the leading pacemaker site. Each leading site was replotted onto the SAN/atrial tissue image.

**Isolation of SAN cardiomyocytes**

SAN tissue was excised by visually cutting along the crista terminalis and the interatrial septum in prewarmed (36 °C) Tyrodes solution. The excised tissue was immersed in a solution containing (in mM):140 NaCl, 5.4 KCl, 0.5 $MgCl_2$, 0.2 $CaCl_2$,1.2 $KH_2PO_4$, 5 HEPES, 5 taurine, 5.5 Dextrose, and 1 mg/mL bovine serum albumin (BSA), adjusted to pH 6.9 using NaOH, for 6 minutes. Digestion was then performed in the same solution containing Liberase™ (350 U/mL; Roche) and Elastase (2 Units/mL, Worthington) for 15–20 min at 36 °C. To terminate the digestion process, the SAN tissue was washed in a modified "Kraftbrühe" (KB) medium containing (in mM): 70 K-glutamate, 25 KCl, 10 $KH_2PO_4$, 20 Taurine, 5 HEPES, 10 K-aspartate, 20 Dextrose, 0.5 EGTA, 2 $MgSO_4$, 5 Creatine, and 1 mg/mL BSA, with the pH adjusted to 7.4 using KOH. Single cells were then obtained by manual agitation of the tissue in KB solution at room temperature for 2 minutes. Ca was gradually reintroduced to a final concentration of 1.8 mM, as we described [2, 3].

**Single Cell Electrophysiology**

Isolated cardiomyocytes were placed in a flow-through optical chamber on the stage of an inverted microscope. The chamber was perfused with 1.8 mM Ca Tyrodes (as described above) or experimental solutions as detailed below (at 20-22°C). Membrane voltage was recorded using an Axopatch 200B patch clamp amplifier (Molecular Devices, Sunnyvale, CA) with signals digitized by a Digidata 1440a A/D converter (Molecular Devices), controlled by pClamp software (version 10.4; Molecular Devices). Patch electrodes were pulled from borosilicate glass (TW150F-3, World Precision Instruments, Sarasota, FL) using a Flaming-Brown horizontal micropipette puller (P-97, Sutter Instruments, Novato, CA). The glass pipettes had an electrical resistance of 1-2 MΩ. Membrane current was filtered at 5 kHz and digitized at 2 kHz.

To measure pacemaker current ($I_f$) using the whole cell technique, the patch pipette internal solution contained (in mM): 128 K-aspartate, 7 KCl, 1 $MgCl_2$, 10 HEPES, 1 $CaCl_2$, 10 EGTA, 6.6 Na-phosphocreatine, 0.1 Na-GTP and 4 Mg-ATP, pH 7.2, as previously used[3, 5]. The bath was a modified Tyrodes solution containing (in mM) 140 NaCl, 5.4 KCl, 1 $MgCl_2$, 10 HEPES, 10 dextrose, 1.8 $CaCl_2$, and 1 $BaCl_2$, with a pH of 7.4.

To record SAN cell action potentials using the perforated patch clamp technique, the internal solution contained (in mM): 130 KCl, 10 NaCl, 10 HEPES, 5 Mg-ATP, 0.05 cAMP, 1 $MgCl_2$, and 5 EGTA (pH adjusted to 7.2 using KOH). EGTA was included to facilitate identification of ruptured patches. The bath solution was the 1.8 mM Ca Tyrodes solution described above. The internal solution was initially drawn into the tip of the pipette using capillary action and then backfilled with the same solution containing 240μM amphotericin B. After obtaining a gigaohm seal, electrical access was typically obtained within 5 minutes. To record spontaneous SAN cell APs, we used the current clamp mode of the Axopatch 200B and recorded voltage changes without applying any electrical stimulation. To evoke APs, we applied a 2 ms current injection, increasing from 0 nA to threshold in 0.1 nA increments.

**RNA Preparation and qPCR**

We used TRIzol™ reagent (Invitrogen, Carlsbad, CA, USA) to extract total RNA from isolated SAN/Atrial tissue preparations, per manufacturer's instructions. After confirming RNA quantity and purity through spectrophotometry, 1 μg of RNA was reverse transcribed into cDNA using the RT2 First Strand Kit (Qiagen, Germantown, MD). The resulting cDNA was mixed with $RT^2$ SYBR Green ROX qPCR Mastermix (Qiagen, Germantown, MD) and $RT^2$ qPCR Primer Assays. PCR was performed on 96-well plates using the 7900 HT Fast-Time PCR system (Applied Biosystems, Life Technologies, Foster City, CA). The following primers were used to detect target transcripts:

| Gene Symbol | Orientation | Sequence (5' -> 3') |
|---|---|---|
| GAPDH | Forward | TCACCACCATGGAGAAGGC |
| | Reverse | GCTAAGCAGTTGGTGGTGCA |
| HCN4 | Forward | GGCGGACACCGCTATCAAA |
| | Reverse | TGCCGAACATCCTTAGGGAGA |
| ZO-1 (Tjp1) | Forward | TTTCAGAGTGGGGAAACCTCC |
| | Reverse | CACTCTTCCTTAGCTGCTGAAC |

We used the comparative $C_T$ method (SDS 2.3 software, Applied Biosystems, Foster City, CA) to evaluate RNA expression levels, using glyceraldehyde 3-phosphate dehydrogenase (GAPDH) as the reference gene. The expression level, calculated as $2^{-\Delta\Delta CT}$, corresponds to fold difference in gene expression, relative to the control sample.

**Statistical analysis**

Statistical analyses were performed using GraphPad Prism 10.4.2 Software. All summary data are presented as mean ± standard error of the mean (SEM). Data normality was assessed using the Shapiro-Wilk test. For normally distributed data, comparisons between two groups were made using an unpaired two-tailed t-test.

For data that were non-normally distributed, we used the two-tailed Mann-Whitney test. In some experiments, a nested t-test or one-way ANOVA was used, as indicated in the accompanying figures. P values < 0.05 were considered statistically significant. We used the following symbols to indicate degree of significance in figures: * p<0.05; ** p<0.01; *** p<0.001; **** p<0.0001

**Supplemental Figure**

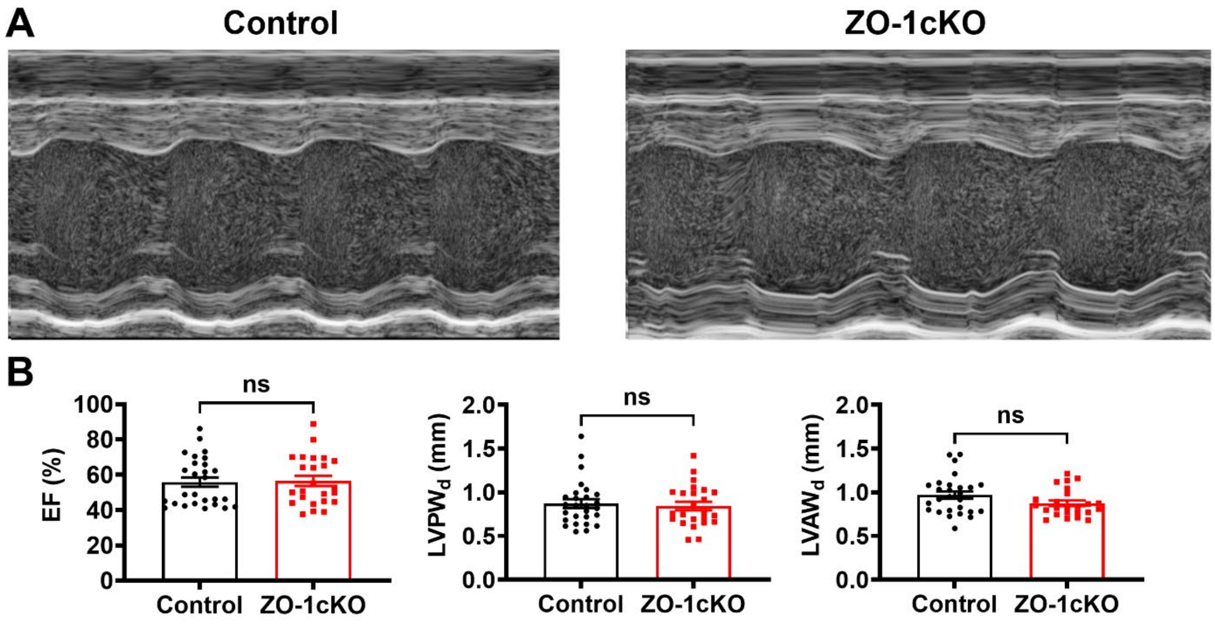


**Figure S1: Normal contractility in ZO-1cKO mice.** A, Representative left ventricular (LV) short axis M-mode views from lightly anesthetized control and ZO-1cKO mice. B, Summary plots show that there were no changes in left ventricular (LV) ejection fraction (EF), LV posterior wall thickness in diastole (LVPWd) and LV anterior wall thickness in diastole (N=24-27 mice in each group). Data were analyzed by Student's t-test and presented as mean ± Standard Error of Mean (SEM).
